# Mapping the adaptive landscape of a major agricultural pathogen reveals evolutionary constraints across heterogeneous environments

**DOI:** 10.1101/2020.07.30.229708

**Authors:** Anik Dutta, Fanny E. Hartmann, Carolina Sardinha Francisco, Bruce A. McDonald, Daniel Croll

**Affiliations:** Plant Pathology, Institute of Integrative Biology, ETH Zurich, Zurich, Switzerland; Ecologie Systématique Evolution, CNRS, Université Paris-Saclay, AgroParisTech, 91400, Orsay, France; Laboratory of Evolutionary Genetics, Institute of Biology, University of Neuchâtel, 2000 Neuchâtel, Switzerland

## Abstract

The adaptive potential of pathogens in novel or heterogeneous environments underpins the risk of disease epidemics. Antagonistic pleiotropy or differential resource allocation among life-history traits can constrain pathogen adaptation. However, we lack understanding how the genetic architecture of individual traits can generate trade-offs. Here, we report a large-scale study based on 145 global strains of the fungal wheat pathogen *Zymoseptoria tritici* from four continents. We measured 50 life-history traits, including virulence and reproduction on 12 different wheat hosts and growth responses to several abiotic stressors. To elucidate the genetic basis of adaptation, we used multi-trait genome-wide association mapping. We show that most traits are governed by polygenic architectures and are highly heritable suggesting that adaptation proceeds mainly through allele frequency shifts at many loci. We identified numerous pleiotropic SNPs with conflicting effects on host colonization and survival in stressful environments. Such genetic constraints are likely limiting the pathogen’s ability to cause host damage and could be exploited for pathogen control. In contrast, adaptation to abiotic stress factors was likely facilitated by synergistic pleiotropy. Our study illustrates how comprehensive mapping of life-history trait architectures across diverse environments allows to predict evolutionary trajectories of pathogens confronted with environmental perturbations.

## Introduction

Adaptation to heterogeneous environments (both biotic and abiotic) is critically important for pathogens to successfully infect and colonize their hosts, disseminate to new hosts and cause epidemics, and survive in the absence of their host. Adaptation to new environments is contingent on genetic variation in life-history traits. Adaptation can be severely constrained in changing environments when pathogen populations carry low genetic diversity or are challenged by trade-offs between advantageous traits (Lannou, 2012; Laines and Barres, 2013). Trade-offs typically arise from differential resource allocation and antagonistic gene actions, including pleiotropy, that inhibit the simultaneous increase in two favorable traits (Anderson and May, 1982; Stearns, 1989; Roff, 2000). Understanding the genetic routes of adaptation and disentangling the relationships among adaptive traits can prove useful for predicting evolutionary responses to selection pressures and designing successful disease management strategies (Trivedi and Wang, 2014; Hughes and Leips, 2017). Despite the likely importance of trade-offs, we largely lack evidence that trade-offs have constrained pathogen adaptation in natural environments. A few studies focusing on specific traits and growth conditions reported mutations conferring thermal trade-offs in *Escherichia coli* (Rodríguez-Verdugo et al. 2014), fitness trade-offs in drug resistance in *Candida albicans* (Hill et al. 2015), and a simultaneous increase in melanin synthesis and virulence in *Aspergillus fumigatus* (Jackson et al. 2009). The mutations underlying synergistic or antagonistic interactions among life-history traits, including virulence and adaptations to environmental stresses, remain largely unknown, hindering our understanding of adaptive landscapes in pathogens.

Trade-offs are defined as phenotypic correlations that prevent organisms from reaching maximum fitness in specific environments and are often influenced by environmental conditions. Trade-offs impose evolutionary constraints if the expression of each trait is governed by the same mutations (Roff, 2000). Pleiotropy occurs when single mutations simultaneously affect the expression of several traits, generating genetic correlations among these traits (Wang et al. 2010; Wagner and Zhang, 2011). Genetic correlations can also arise from linkage disequilibrium among mutations, but such correlations are less stable over generations as recombination tends to break down the association (Roff, 2007; Saltz et al. 2017; Hughes and Leips, 2017). Investigation of pleiotropic effects has largely focused on a small number of traits in different species (Scarcelli et al. 2007; Lendenmann et al 2015; Fletcher et al. 2015; Durmaz et al. 2019; Chen and Zhang, 2020). To elucidate the constraints on adaptation in pathogens will require a genome-wide perspective that can consider trade-offs and pleiotropy across a broad spectrum of fitness-relevant life-history traits. It is particularly challenging to identify trade-offs and pleiotropy when the genetic architecture of the studied traits is dominated by numerous small effect loci (*i.e.* polygenic traits), yet the majority of phenotypes affecting fitness show this genetic architecture (Hill and Zhang, 2012). We aimed to establish a fine-scale map of genetic correlations to unravel constraints on trait evolution in host and non-host environments.

*Zymoseptoria tritici* is a major fungal pathogen that poses a significant threat to global wheat production by causing septoria tritici blotch (STB) disease (Fones and Gurr, 2015; Torriani et al. 2015). The life cycle of the pathogen is complex, including niche adaptations that enable growth and survival on host tissue across a wide range of temperatures, asexual reproduction that enables short distance dispersal and the production of survival propagules at the end of the growing season (Croll and McDonald, 2017). The pathogen causes necrotic lesions on infected leaves and reproduces asexually with fruiting bodies called pycnidia that form within lesions and produce spores that are dispersed by rain splash. Lesion development and pycnidia production are core aspects of pathogen virulence and asexual reproduction, respectively. Lesion development and pycnidia production show strong correlations at the phenotypic level that are consistent with trade-offs (Dutta et al. 2020). Later in the infection cycle, the pathogen forms sexual ascospores that are dispersed over long distances by wind. Across the global distribution range, populations vary in their degree of thermal adaptation and fungicide resistance as a result of differences in temperature and fungicide exposure (Zhan et al. 2006; Zhan and McDonald, 2011). The pathogen produces variable levels of melanin under stress conditions and melanin production shows negative correlations with growth rates and fungicide susceptibility (Lendenmann et al. 2015; Krishnan et al. 2018). Under high temperature, the pathogen also produces resistant survival structures called chlamydospores (Francisco et al. 2019). As for most pathogens, the genetic architecture underlying life history traits, including virulence, reproduction and stress resistance remains largely unknown. The *Z. tritici* model offers powerful experimental approaches to map phenotypic trait architectures using genome-wide association mapping (GWAS; Sánchez-Vallet et al. 2018), shows high heritable trait variation in populations (Hartmann et al. 2017) and extensive genomic resources are available (Goodwin et al. 2011; Badet et al. 2020) that can enable the precise localization of adaptive loci.

Here, we perform a large-scale, multi-trait GWAS in a global collection of 145 *Z. tritici* isolates originating from five field populations spanning four continents. We map adaptive loci for a total of 50 phenotypic traits, including virulence and reproduction on 12 distinct host genotypes covering both landrace diversity and elite cultivars. Adaptation to environmental factors was assessed for different temperatures and fungicide concentrations. Combining information about adaptive loci across the genome, we analyze the extent of genetic correlations among classes of phenotypic traits to identify major trade-offs. By mapping the adaptive landscape of constraints and facilitation governing phenotypic trait evolution, we generate a comprehensive view on the genetic underpinnings of pathogen evolution.

## Results

### Extensive trait variability in a highly polymorphic pathogen

We analyzed the expression of life-history and physiological traits in a panel of *Z. tritici* isolates exposed to different environmental conditions. The collection spans the distribution range of the pathogen including the Middle East (the pathogen center of origin), Europe, North America and Australia (**Figure 1A**). To assess pathogenicity trait variation, we infected 12 distinct wheat cultivars and measured the capacity to cause damage and reproduce on the host. We also assessed growth and colony morphology under culture conditions applying a variety of stress factors (**Figure 1B**). Isolates showed a high degree of phenotypic diversity across all measured traits (**Figure 1C & Supplementary Table S2**). Trait expression was largely quantitative and almost all traits showed substantial variation within each field population. While there was no obvious geographic clustering of isolate phenotypes according to geographic origin (*i.e.* population structure), we found considerable differences among populations for mean trait values (**Figure 1D**). The Israeli population showed the highest host specialization for reproduction and also the highest percentage of isolates that formed resistance structures (*i.e.* chlamydospores) under stress (Francisco et al. 2020). The Swiss population showed the highest fungicide resistance (*P<0.0001*) consistent with the earlier and more pervasive application of azole fungicides in Europe compared to other regions (Zhan et al. 2006). The extensive variation in phenotypic traits across host and non-host environments suggests that these traits are governed by complex genetic architectures. Hence, identifying the loci underpinning the trait variation across genotypes can provide insights into the evolutionary trajectory of these pathogen populations.

**Figure 1.**
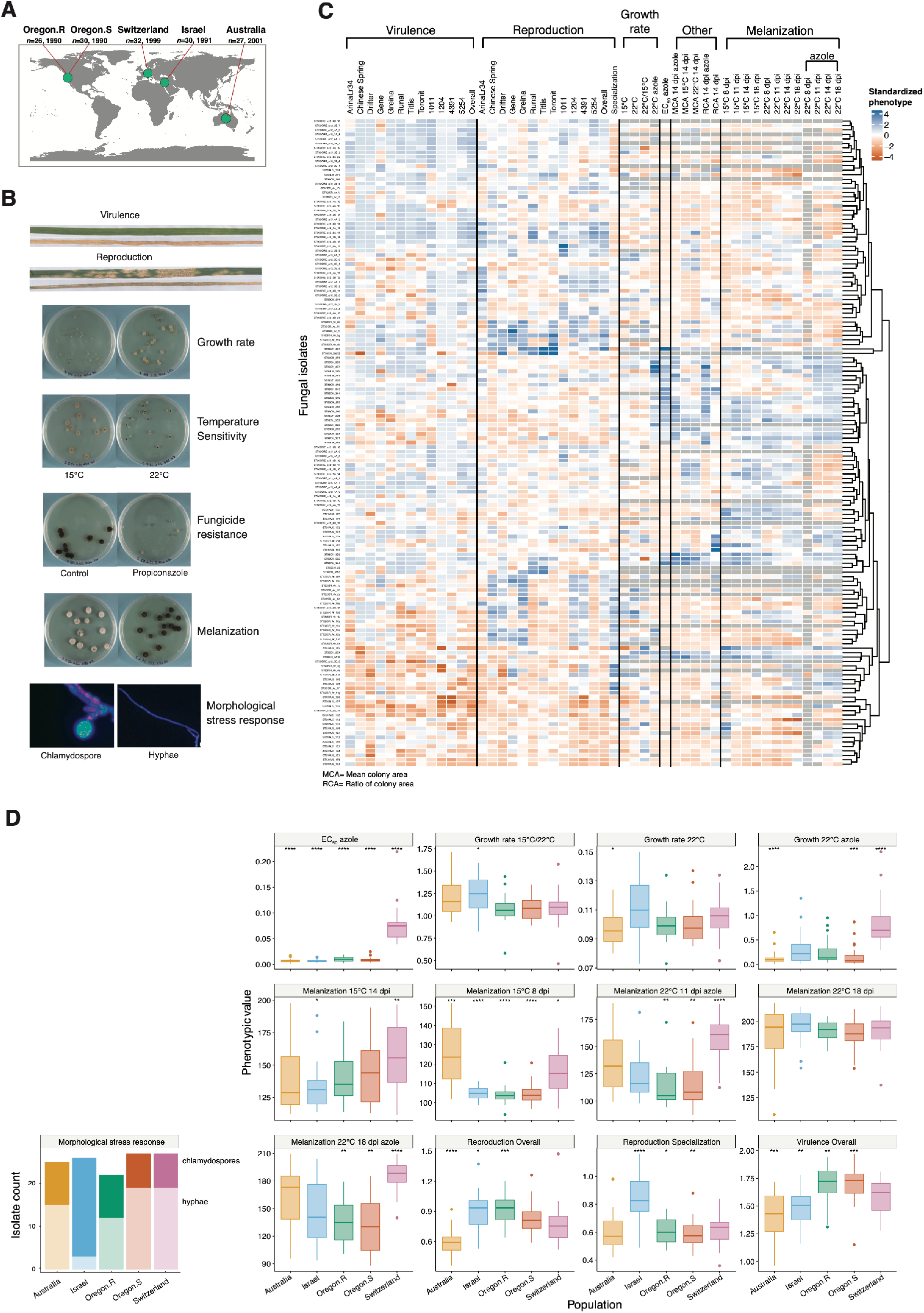
Geographic origin of populations and phenotypic diversity of *Zymoseptoria tritici*. (A) World map showing the field locations of the five pathogen populations (*n* = 145 isolates). The two populations in Oregon (USA) were sampled from wheat cultivars Madsen (resistant; Oregon.R) and Stephens (susceptible; Oregon.S). (B) Major categories of life-history traits observed in *Z. tritici* on the host and in the non-host environments (virulence – amount of necrotic lesion on the leaf; reproduction – pycnidia formation within lesions). (C) Heatmap showing phenotypic diversity of 49 traits using standardized phenotypic trait values. Pathogen virulence was assessed by the percentage of necrotic lesions on wheat leaves. Reproduction specialization was defined by the adjusted coefficient of variation of means across all 12 wheat hosts. Melanization was expressed on a grayscale ranging from 0 (white) to 255 (black). Dendrogram branches correspond to Euclidean distances for phenotypic trait values. **(D)** Phenotypic trait distribution in different environments among the five populations. The binary morphological stress response trait is shown separately.

### Trait heritability for host adaptation and survival in the environment

We estimated SNP-based heritability (*h*^*2*^_*snp*_) using relatedness-based restricted maximum likelihood to partition the observed phenotypic variation. The estimated *h*^*2*^_*snp*_ for virulence ranged from 0 to 0.59 (standard error, SE=0.14) and for reproduction from 0.43 (SE=0.15) to 0.91 (SE=0.03; **Figure 2A**). The higher mean *h*^*2*^_*snp*_ for reproduction (0.72, SE=0.10) compared to the mean for virulence (0.47, SE=0.15) suggests that the short-term response to selection is likely to be faster for reproduction. The *h*^*2*^_*snp*_ for growth, fungicide resistance, morphological stress response and thermal tolerance traits ranged from 0 to 0.99 with fungicide resistance traits showing the highest heritability (**Figure 2B**). This is consistent with previous findings of target gene mutations playing a dominant role in fungicide resistance (Cools et al. 2011; McDonald et al. 2019).

**Figure 2.**
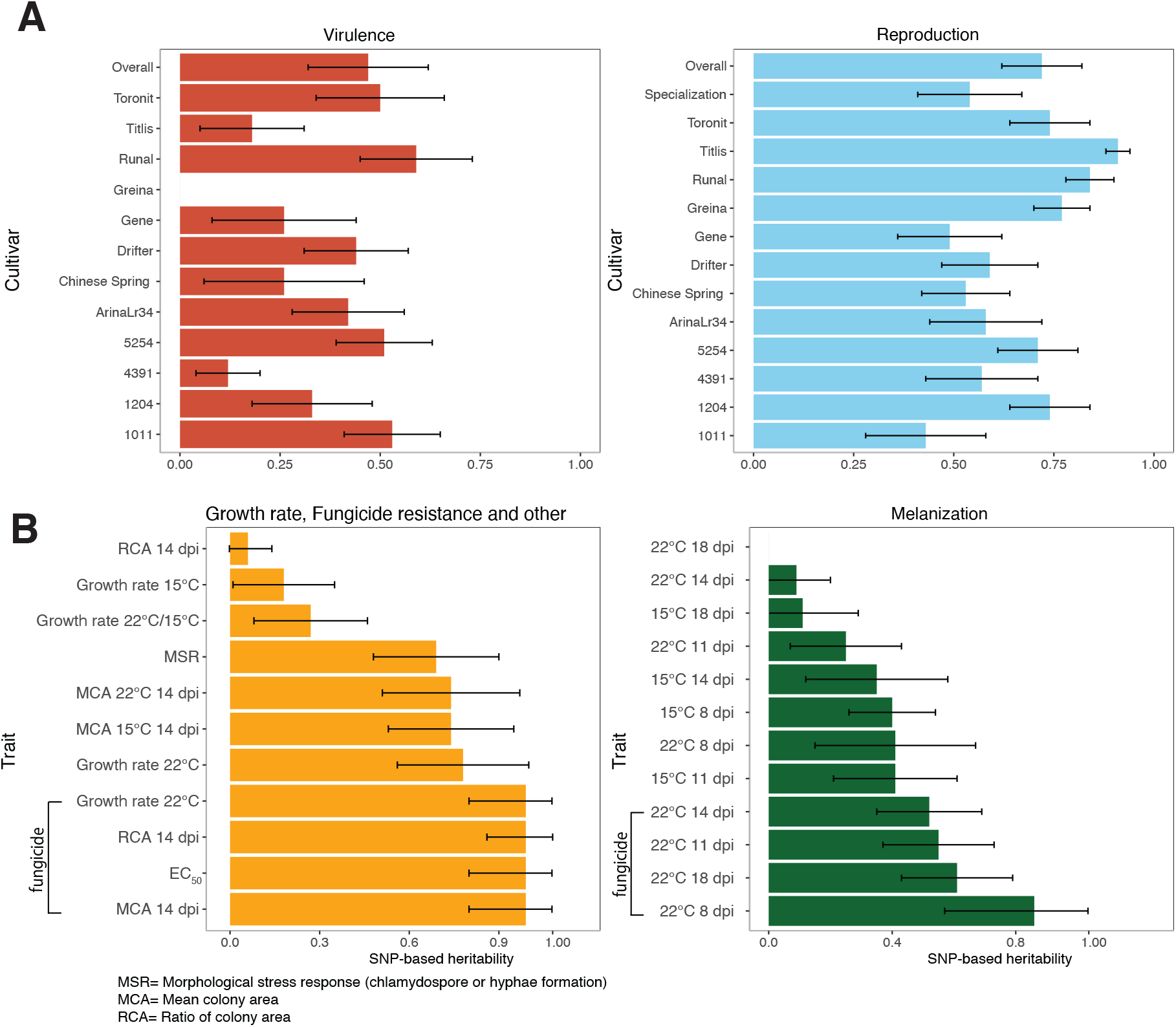
Phenotypic variance explained by common genetic variants. (A) Pathogen virulence (amount of necrotic lesion area) and reproduction (pycnidia density within the lesion area) were measured on 12 wheat hosts. (B) Growth rate in absence of the host, fungicide resistance, melanization and morphological stress response (chlamydospore or mycelium production). SNP-based heritabilities (*h*^*2*^_*snp*_) were estimated by following a GREML approach. Error bars indicate standard errors.

We performed GWAS on each trait to identify genetic variants associated with trait expression across different environments. For the majority of the traits, including virulence, reproduction and fungicide resistance, we detected between 18-1422 significantly associated SNPs at a 10% false discovery rate (FDR) threshold. We identified no phenotype-genotype association at the more stringent Bonferroni threshold (α = 0.05). This suggests a polygenic architecture with many loci of small effect underlying each trait. In addition, we identified significant SNPs which were shared among multiple traits **(Figure 3A & Supplementary Table S3)**. We detected 249 SNPs shared among fungicide resistance traits including mean colony area at 14 dpi at 22°C and the ratio of colony area at 14 dpi in presence of fungicides **(Figure 3B)**. Importantly, we detected four shared SNPs associated with virulence on the landraces 1204 and 4391, as well as 11 SNPs associated with reproduction on the elite cultivars Greina, Titlis and Toronit **(Figure 3B)**. The discovery of shared SNPs underlying variation in pathogenicity among multiple cultivars and with traits expressed outside of the host suggests that host adaptation is possibly constrained by pleiotropic effects.

**Figure 3.**
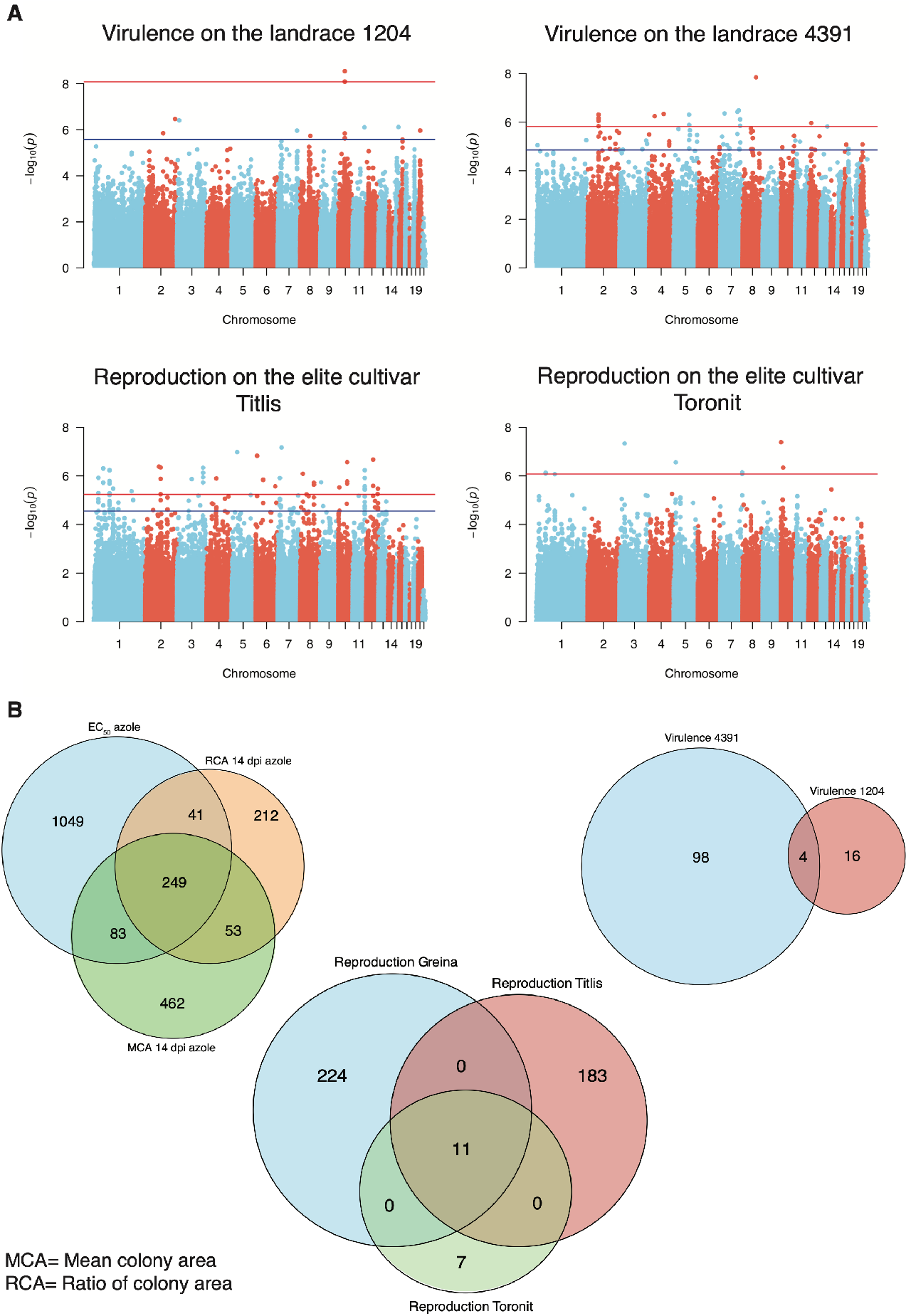
Genome-wide association mapping (GWAS) for 50 traits measured in various host and non-host environments. (A) Manhattan plots showing SNP marker association *p*-values for four individual traits. The blue and red lines indicate the significant thresholds for false discovery rates (FDR) at 10% and 5%, respectively. (B) Venn diagrams showing the number of SNPs significantly associated with multiple traits (FDR 10%). Virulence (amount of necrotic lesion area) and reproduction (pycnidia density within the lesion area) were measured on 12 different wheat hosts.

### Trait correlations and complex genetic control in variable environments

Phenotypic correlations can arise from both shared genetic architectures and confounding effects of population substructure (*i.e.* linkage disequilibrium). Here, we focus on genetic correlations estimated from GWAS using a mixed-effect model to account for genetic relatedness among individuals. To quantify the overlap in genetic architectures between traits, we analyzed genetic correlations using the estimated allelic effects at all SNPs derived from individual GWAS (**Figure 4**). We also analyzed phenotypic correlations (*r*_*p*_) using standardized phenotypic trait values. Both genetic correlations of allelic effects between traits (*r*_*g*_= −0.81 to 0.89, 0.91 ≤ *P* < 4.44e^−16^) and phenotypic correlations (*r*_*p*_= −0.83 to 0.91, 0.99 ≤ *P* < 6.6e^−16^) were highly variable (**Supplementary Table S4**). We found a significant genetic trade-off between specialization in reproduction and overall virulence (*r*_*g*_= −0.67, *P* < 4.44e^−16^). The significant positive genetic correlations (*r*_*g*_= 0.03 to 0.34, < 4.44e^−16^) between morphological stress response (*i.e.* chlamydospore formation or hyphal growth) and colony melanization in multiple environments suggests a shared genetic control for these traits. Furthermore, fungicide resistance showed a highly significant negative genetic correlation with virulence (*r*_*g*_= −0.17, *P* < 6.6e^−16^) and reproduction (*r*_*g*_= −0.23, *P* < 6.6e^−16^) on the cultivar Gene, reproduction on the landrace 1204 (*r*_*g*_= −0.22, *P* < 6.6e^−16^) and the degree of reproduction specialization (*r*_*g*_= −0.24, *P* < 6.6e^−16^). Overall colony growth rates at 22°C had mostly very low genetic and phenotypic correlations with pathogenicity traits. This suggests that mutations favoring colony growth may not favor overall plant colonization. Overall, the observed genetic correlations indicate extensive pleiotropy among genes controlling major phenotypic traits of the pathogen.

**Figure 4.**
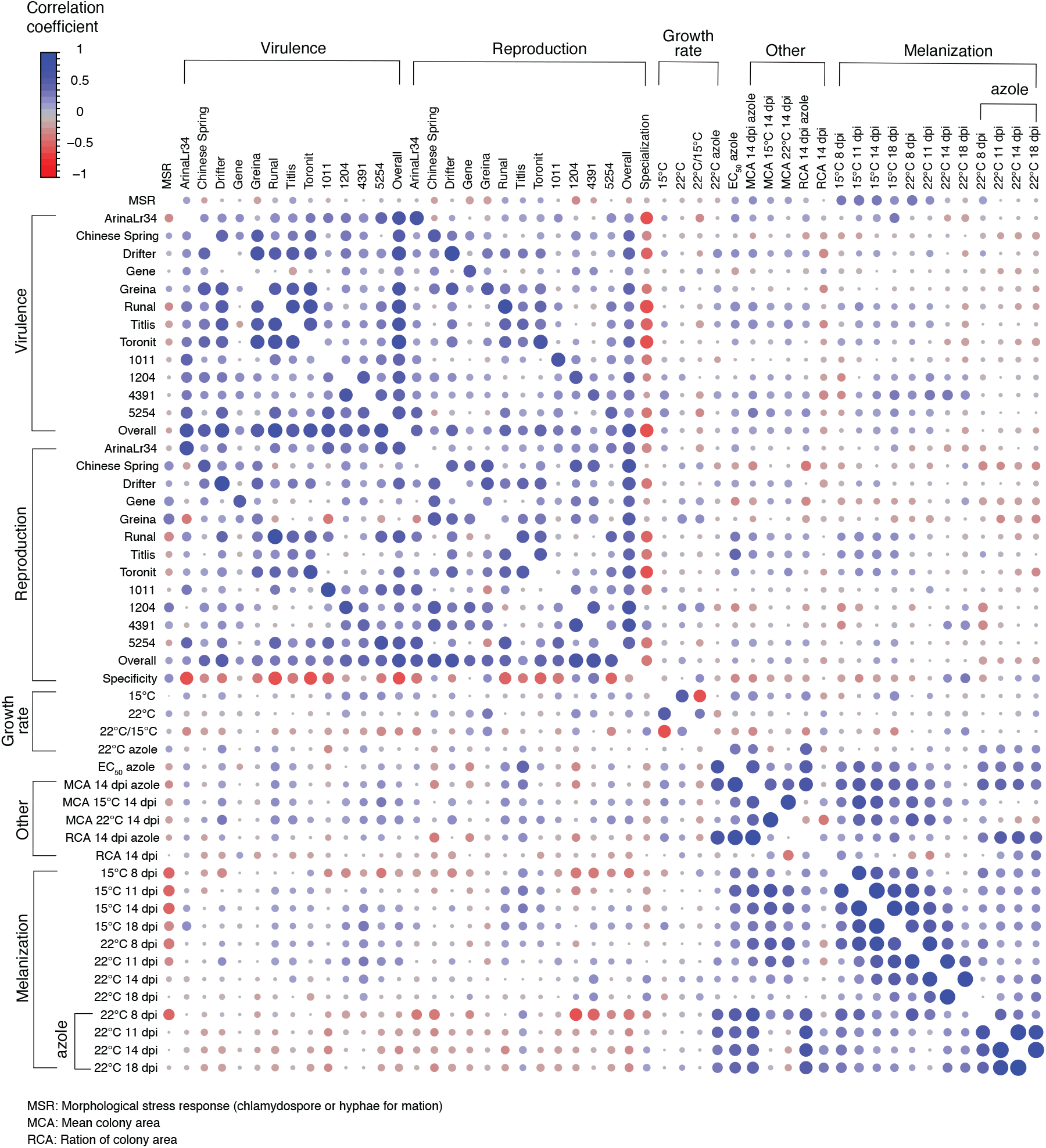
Genetic (upper diagonal) and phenotypic (lower diagonal) correlations among 50 traits measured in various host and non-host environments. Genetic correlations were estimated using the GWAS-derived allelic effects. Phenotypic correlations were estimated using standardized phenotypic values. Blue and red colors indicate positive and negative correlation coefficients, respectively. The circle size and color intensities are proportional to the correlation coefficient. Pathogen virulence (amount of necrotic lesion area) and reproduction (pycnidia density within the lesion area) were measured on 12 different wheat hosts.

We quantified the difference between phenotypic and genetic correlations for each pair of traits. Focusing on the most extreme differences (**Figure 5A**), we found an excess of 0.75 in the genetic correlation for the morphological stress response and melanization (at 15°C 11 dpi; *r*_*g*_= 0.34, *r*_*p*_= −0.41) and an excess of 0.57 in the phenotypic correlation for the morphological stress response and reproduction on the cultivar Greina (*r*_*g*_= −0.19, *r*_*p*_ = 0.38). Given this discordance, we therefore used only genetic correlation coefficients for the subsequent analyses. The average *r*_*g*_ within each trait and between pairs of traits showed the relative overlap of loci having similar effects in different environments (**Figure 5B**). The average *r*_*g*_ for virulence (0.31±0.02) and reproduction (0.19±0.02) was higher than for growth rate (0.005±0.15). This indicates that traits related to plant colonization exhibit stronger correlations among different hosts than colony growth outside of the host. The highest average genetic correlation was found for melanization (0.35±0.02) followed by fungicide resistance and associated traits for colony area (other; 0.32±0.11), virulence and melanization and fungicide resistance (0.28±0.04). Our findings show that the genetic architecture of melanization and fungicide resistance across environments is strongly overlapping.

**Figure 5.**
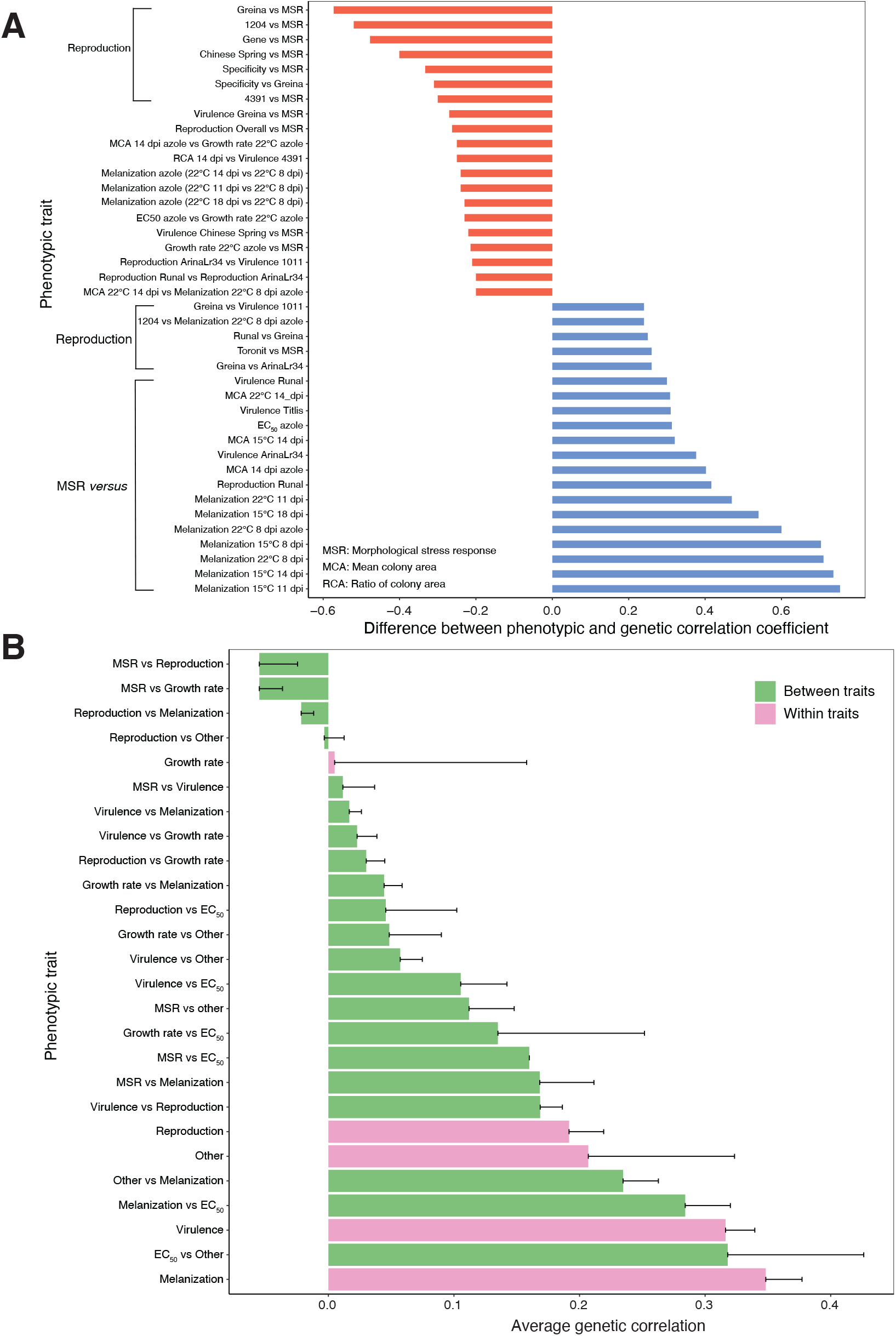
Discordance between genetic and phenotypic correlations. (A) Bar plot showing the difference between genetic and phenotypic correlation coefficients. A total of 20 trait pairs from each tail of the distribution (*n* = 1225 pairings) were selected for visualization. (B) Bar plot showing the average genetic correlation (relative degree of overlapping loci) within and between major trait categories. Pathogen virulence (amount of necrotic lesion area) and reproduction (pycnidia density within the lesion area) were measured on 12 diverse wheat hosts. Error bars indicate standard errors.

The host genetic background shows strong interactions with loci contributing to pathogenicity. Hence, we analyzed how strongly pathogenicity traits correlate between hosts (**Figure 6A-C**). We found that the average *r*_*g*_ for virulence and reproduction on the elite cultivar Gene (0.06±0.03, 0.16±0.04, respectively) and the landrace 1011 (0.17±0.03, 0.12±0.03, respectively) were particularly low compared to the other cultivars. This indicates that the genetic control of pathogenicity on those two hosts is the most differentiated compared to the other hosts. The negative average *r*_*g*_ for reproduction specialization (−0.22±0.06) is in contrast to the positive overall average *r*_*g*_ for virulence (0.56±0.03) and reproduction (0.52±0.02) (**Figure 6D**). This suggests antagonistic pleiotropy in the genetic control of host specialization. We further investigated how genetic control for pathogenicity on different hosts overlaps with genetic control of colony growth, temperature sensitivity and fungicide resistance (**Figure 6E-H & Supplementary Figure 1**). We found a positive average *r*_*g*_ for all the above traits on the elite cultivars Titlis and Runal. This suggests synergistic pleiotropy of pathogenicity and performance in absence of the host (*i.e.* stress response, thermal sensitivity and growth). Pathogenicity on the other hosts shows a wide range of positive and negative average *r*_*g*_ between different traits. Hence, SNPs associated with increased virulence and reproduction on specific hosts can have antagonistic effects on other traits such as growth or melanization. This suggests that a complex mix of antagonistic and synergistic pleiotropy underlies the evolution of pathogenicity.

**Figure 6.**
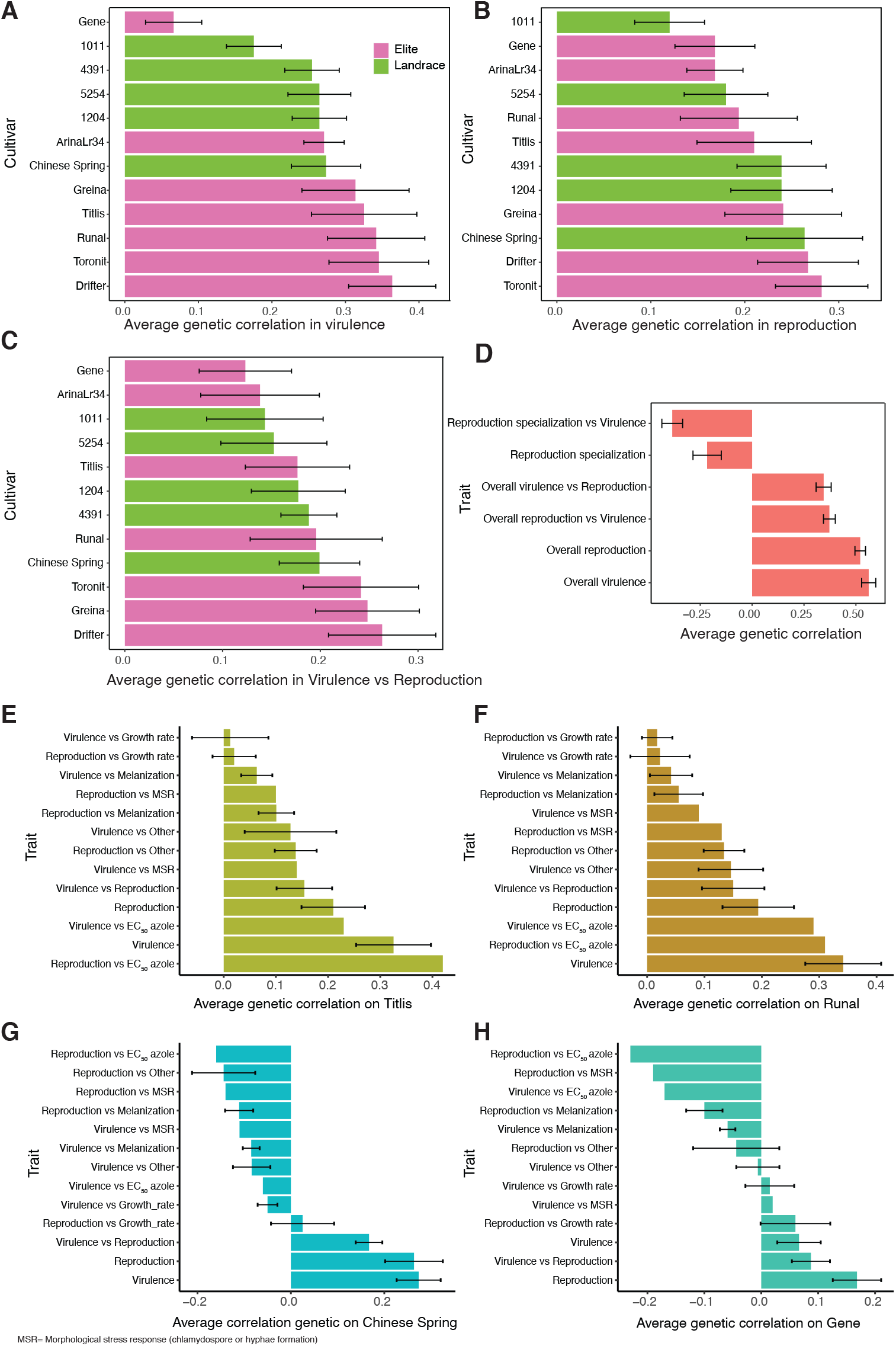
Average genetic correlation among host adaptation traits and environmental stress traits. Bar plots represent shared adaptive loci inferred from average genetic correlation coefficients for (A) pathogen virulence (amount of necrotic lesion area), (B) reproduction (pycnidia density within the lesion area), (C) between virulence and reproduction on each of the 12 wheat hosts, (D) overall virulence and reproduction combined over 12 hosts and host specialization. Host specialization is represented as the coefficient of variation of means across 12 hosts. (E-H) Bar plots showing the relative degree of interaction between the genetic control of host and non-host traits on each of the 12 hosts. Only four hosts are shown and the eight remaining hosts are shown in Supplementary Figure 1. Melanization was expressed on a grayscale ranging from 0 (white) to 255 (black). Error bars indicate standard errors.

## Discussion

A mechanistic view of the adaptive landscape across the life cycle is of fundamental importance to understand pathogen evolution (Hartmann et al. 2019). Yet, we lack fundamental insights into the underlying genetic architecture and how adaptation may be constrained. Here, we report the first study that maps a broad range of adaptive traits in a pathogen using a large-scale multi-trait GWAS. We show that the expression of most life-history traits is highly variable at the population level and has a largely polygenic basis. We found strong evidence of genetic trade-offs and facilitation among the life-history traits.

### Polygenic architecture of life-history trait expression

We show that a substantial proportion of the overall phenotypic variation is explained by common variants (*i.e.* SNPs with minor allele frequency > 0.05), consistent with polygenic trait architectures with many loci of small individual effect (Yang et al., 2010, 2017; Korte and Farlow, 2013; Slate, 2013; Wood et al., 2014). A polygenic architecture for major life history traits means that adaptation across the diverse environments on and off the host likely proceeds via subtle shifts in allele frequencies across many loci (Höllinger et al. 2019). The polygenic nature of most traits was further supported by the identification of loci controlling traits measured in multiple host and non-host environments. We found that heritability was higher for pathogen reproduction than pathogen virulence (*i.e.* damage to the host). This may be explained by more differentiated host exploitation strategies for reproduction contributing to the maintenance of variation within pathogen populations. Higher heritability for pathogen reproduction on hosts means a potentially faster response to selection and ultimately host adaptation. However, selection on reproduction should also impact virulence and transmission through correlated responses (Anderson and May, 1982; Pettay et al., 2005; Leggett et al., 2013; Dutta et al., 2020). Variation in heritability for fungicide resistance, thermal adaptability, and stress response could be attributed to heterogeneous selection pressures across the worldwide sampling range. Genetic variation for traits expressed outside of the host is thought to facilitate pathogen adaptation to abiotic stress. Maintenance of such variation is particularly relevant as pathogen populations face potential bottlenecks in-between the annual wheat growing seasons (McDonald and Linde, 2002; Zhan et al., 2005). A relevant consideration is whether the homogenization of agricultural landscapes impacts pathogen evolution (Stukenbrock and McDonald, 2016). Homogenization of the environment can increase directional selection pressure and erode heritable variation for life history traits. How *Z. tritici* retains such high levels of adaptive genetic variation along environmental gradients remains unclear. A key factor maintaining variation may be genetic trade-offs among life history traits constraining directional selection (Reznick, 1992; Roff, 1992).

### The adaptive landscape is shaped by extensive genome-wide pleiotropy

Adaptation to complex environments can be severely constrained by genetic trade-offs, but populations can also experience rapid adaptation through correlated responses to selection if genetic correlations are positive (Stearns 1989; Roff 2000; Sgrò and Hoffmann, 2004; McGee et al., 2016). We show that adaptation of *Z. tritici* is both constrained and facilitated by genetic correlations that vary according to the life-history traits and the environment. Unlike phenotypic correlations, genetic correlations estimated using mixed models are robust to confounding effects of population structure and directly reflect the genetic contributions to adaptive trait co-variation (Sullivan et al., 2015; van Rheenen et al., 2017). Given the largely polygenic architecture of most life history traits, the observed genetic correlations are most likely caused by pleiotropic effects of numerous small effect loci. Alternatively, genetic correlations could be controlled by nearby loci in linkage disequilibrium. However, such genetic correlations should be transient and broken down rapidly by the high recombination rate of *Z. tritici* (Zhan et al., 2003, 2005; Croll et al., 2015). Overall, the magnitude of trade-offs and facilitation varied considerably among biotic and abiotic environments, and life cycle stages. Precise knowledge of genetic correlations can identify trade-offs and through this help to predict the evolutionary trajectories of pathogen populations.

A major finding on the evolution of pathogenicity is the positive correlations between virulence and reproduction. Such correlations could stem from convergent evolution selecting for similar synergistic mutations in shared resistant backgrounds of cultivated wheat worldwide (Brown et al., 2015; Croll and McDonald, 2017). The considerable overlap in shared adaptive loci likely facilitates rapid adaptation to diverse host backgrounds. Adaptation to resistant hosts such as Gene or the landrace 1011 very likely imposed selection pressure on pathogen populations to evolve a distinct, host-specific arsenal of virulence factors. Consistent with theoretical predictions (Kassen, 2002; Legros and Koella, 2010), our study suggests that adaptation to novel hosts will be constrained due to strong antagonistic effects of host specialization for reproduction. Furthermore, adaptation to new abiotic environments is likely to be similarly constrained. Host or niche specialization is a driver of pathogen local adaptation, yet antagonistic effects can result in pathogen maladaptation when faced with nonoptimal environments (Croll and McDonald, 2017). In contrast to other pathosystems (e.g. Thrall et al., 2005; Lee at al., 2015), genetic control of *Z. tritici* growth outside of the host was largely independent and distinct from traits promoting host infection, with only few loci showing pleiotropic effects. Nevertheless, on some host backgrounds, the ability to cause disease could be constrained by the pleiotropic effects of stress-related traits such as fungicide resistance or melanization. Genetic constraints governing growth in the absence of the host and pathogenicity could be a major impediment for pathogen populations to maximize fitness on hosts. This would be due to the fact that fitness outside the host is affected by correlated responses to selection on traits expressed on the host. The finding that *Z. tritici* populations carry variants with negative pleiotropic effects for infection despite apparent costs is intriguing. Variants underlying strong stress-related trait expression variation should be highly selectable and are likely underpinning survival under harsh climate as shown by strong synergistic pleiotropic effects. Therefore, maladaptive trait expression on the host could well be maintained in pathogen populations if the underlying variants are favored under harsh climatic conditions (Wright, 1984; Foster et al., 2004; Frénoy et al., 2013).

### Conclusion

Our study reveals remarkable complexity in pleiotropic effects across the life cycle in a major eukaryotic plant pathogen. Furthermore, adaptation to host and non-host environments has a largely polygenic basis. Our study demonstrates the power of conducting a multi-trait GWAS across a wide range of environmental conditions. We show that trade-offs are not ubiquitous, but rather facilitation (*i.e.* synergistic pleiotropy) plays an important role in determining the evolutionary potential of pathogen populations across environments. By leveraging information on facilitation and trade-offs, the response of pathogens to changing environments becomes more predictable. Furthermore, knowledge on trade-offs can be exploited in innovative disease management models that exploit evolutionary weaknesses of pathogens. For example, trade-offs between pathogen virulence on specific cultivars and fungicide resistance could be exploited by planting specific wheat genotypes in areas where fungicide resistance emergence becomes a risk. A clear understanding of facilitation and correlated responses to selection can also be applied to prevent catastrophic breakdowns in resistance. The antagonisms among life-history traits on different hosts can inform crop resistance management by spatially or temporally diversifying agricultural fields with different hosts according to the prevalent environmental conditions. By creating complex but specifically designed host environments, pathogen populations could be constrained in their ability to cause serious host damage. Retracing specific biochemical and developmental pathways underpinning pleiotropic effects is greatly facilitated by the multi-trait genome-wide mapping approach and will provide a deeper insight into principles governing pathogen adaptation in heterogeneous environments.

## Materials and Methods

### Fungal material

We used a panel of 145 *Z. tritici* strains (Supplementary Table S1) sampled from single wheat fields, each planted with a single wheat cultivar, in Australia (*n=27*), Israel (*n=30*), two nearby field sites in Switzerland (*n=2* and *n=30*) and USA (Oregon.R, *n=26*; Oregon.S, *n=30*) between 1990 and 2001 (Zhan et al. 2005). The populations Oregon.R and Oregon.S were sampled from the moderately resistant wheat cultivar Madsen and the susceptible cultivar Stephens, respectively, planted in the same field. Field populations were screened for clones and the analyzed set contains only genetically distinct isolates (Linde et al. 2002; Zhan et al. 2005). Since collection, spores of each strain were stored and maintained in 50% glycerol or anhydrous silica at −80°C.

### Whole-genome sequencing and SNP calling

In addition to the already deposited Illumina whole genome sequencing data for 130 isolates under the BioProject ID PRJNA327615 (Hartmann et al. 2017), we generated raw sequence data for the remaining 15 isolates. Briefly, high-quality genomic DNA from each isolate was extracted following the DNeasy Plant Mini Kits (Qiagen) protocol and paired-end sequencing of 100 bp with an approximately 500 bp insert size was performed using the Illumina HiSeq2000 platform. The newly generated raw sequences were deposited under the BioProject PRJNA327615. We used Trimmomatic v.0.36 (Bolger et al. 2014) to trim low-quality sequencing reads and remove contaminated adapters in each isolate. The refined sequences were aligned to the *Z. tritici* reference genome IPO323 (Goodwin et al. 2011) using Bowtie2 v.2.3.3 (Langmead et al. 2009) and duplicate sequences were removed using MarkDuplicates in Picard tools v.1.118 (http://broadinstitute.github.io/picard). We used Genome Analysis Toolkit (GATK) v.4.0.1.2 (McKenna et al. 2010) for single nucleotide polymorphism (SNP) calling and variant filtration. The GATK HaplotypeCaller was used on each isolate with the command-emitRefConfidence GVCF and -sample_ploidy 1. Joint variant calls were performed using GenotypeGVCFs on a merged gvcf variant file with the option-maxAltAlleles 2. We used VariantFiltration and SNPs were removed if any of the following filter conditions applied: QUAL<250; QD<20.0; MQ<30.0; −2 > BaseQRankSum > 2; −2 > MQRankSum > 2; −2 > ReadPosRankSum > 2; FS>0.1. The final dataset was obtained by filtering with a genotyping call rate of 80% and minor allele frequency (MAF) >5%, which resulted in 716’619 biallelic SNPs across all 21 chromosomes.

### In planta phenotyping

We used phenotypic data on pathogen virulence and reproduction obtained by Dutta et al. (2020) on a panel of 12 genetically different wheat populations including five landraces (Chinese Spring, 1011, 1204, 4391, and 5254), six commercial varieties (Drifter, Gene, Greina, Runal, Titlis, Toronit) and a back-cross line (ArinaLr34). ArinaLr34 carries the wheat leaf rust resistance gene Lr34, which provides resistance to wheat stripe rust (McIntosh, 1992) and powdery mildew (Spielmeyer et al. 2005) but was not previously tested for resistance to *Z. tritici*. The 1011, 1204, 4391, and 5254 landraces were selected from the Swiss National Gene Bank (www.bdn.ch). Six pots with three seeds of each cultivar were sown per pot and placed on a tray in a 2×3 array. The experiment was divided into two phases (6 cultivars at a time) due to space limitations. The trays were kept at 22°C (day) and 18°C (night) with 70% relative humidity (RH) and a 16-h photoperiod in a greenhouse chamber. Each tray containing two-week-old seedlings of six cultivars was inoculated uniformly with each isolate using an airbrush spray gun until run-off. Three independent inoculations were performed to generate three biological replications in different greenhouse chambers in both phases. All leaves from each host-by-isolate combination were collected on the same day between 19-26 days post inoculation (dpi) and analyzed using automated image analysis (AIA; Karisto et al. 2018). The AIA provided quantitative estimates of the necrotic lesion area and pycnidia density within the lesion area. The lesion area and pycnidia density were used as proxies for virulence and reproduction, respectively. The phenotyping procedures are described in more detail in Dutta et al. (2020).

### In vitro phenotyping

Fungal colony growth rate (mm per day), temperature sensitivity, mean colony area, fungicide resistance, and melanization in the presence or absence of fungicide were measured *in-vitro*. Data on the morphological stress response (*i.e.* formation of chlamydospores or hyphae) was obtained from Francisco et al. (2020). The methods used for the *in vitro* phenotyping were adopted from earlier studies (Lendenmann et al. 2014, 2015, 2016; Mohd-Assaad et al. 2016). Briefly, each isolate was regenerated from long-term storage conditions and grown on Petri dishes containing yeast malt sucrose agar (4 gl^− 1^ yeast extract, 4 gl^− 1^ malt extract, 4 gl^− 1^ sucrose, 50 mgl^− 1^ kanamycin) for four to five days at 18°C. Blastospore suspensions were collected from each plate by adding 0.6 ml of sterile water and diluted to a final concentration of 200 spores/ml using KOVA counting slides (Hycor Biomedical, Inc., Garden Grove, CA, USA). For each isolate, a 500 μl spore suspension was spread on Petri plates containing potato dextrose agar (PDA, 4 g l^− 1^ potato starch, 20 g l ^− 1^ dextrose, 15 g l^− 1^ agar) using a sterile glass rod. Plates were kept at 15°C and 22°C at 70% RH. Each plate was photographed at 8, 11 and 14 dpi with a digital camera.

The digital images of each plate from five technical replications were used for AIA in ImageJ following the scripts by Lendenmann et al. (2014) to obtain colony area information for each isolate. Estimates are based on an average of 45 spore colonies. The mean colony radii taking the square-root of mean colony area (*√(mean colony area/Π*)) were estimated and fitted over the three time points following a generalized linear model to obtain growth rates (mm/day) at 15°C and 22°C. Temperature and fungicide sensitivity of each isolate was calculated using the growth rate ratio between 15°C and 22°C, between 22°C and growth rate at 22°C on PDA amended with propiconazole (Syngenta, Basel, Switzerland; 0.05 ppm), respectively. In addition, we used mean colony area measured for each isolate at 14 dpi on PDA at 15°C (MCA_15°C_14_dpi), 22°C (MCA_22°C_14_dpi), and 22°C amended with 0.05 ppm propiconazole (MCA_14_dpi_azole), to estimate the ratio of colony area in cold (RCA_14_dpi) and fungicide environments (RCA_14_dpi_azole) compared to the control environment.

Fungicide resistance assays were carried out using propiconazole on microtiter plates. Growth inhibition was tested by growing spores adjusted to a spore concentration of 2.5 × 10^4^ spores/ml) in Sabouraud-dextrose liquid medium with the following concentrations of propiconazole: 0.00006, 0.00017, 0.0051, 0.0086, 0.015, 0.025, 0.042, 0.072, 0.20, 0.55, 1.5 mg/L and a control without fungicide. Each microtiter well was filled with 100 μl of medium at a given concentration of fungicide and 100 μl of spore suspension using the above spore concentration. Five technical replicates were used for each isolate. The plates were sealed and incubated in the dark for four days at 22°C with 80% relative humidity. Fungal growth was measured with an Elisa plate reader (MR5000, Dynatech) by examining the optical density (OD) at 605 nm wave length. The growth inhibition at different fungicide concentrations was used to estimate the dose-response curves. The dose-response curves were used to estimate the EC_50_ value for each isolate using the drc v.3.0-1package (Ritz et al., 2015) in the R-studio (R Development Core Team, 2019).

We estimated melanization of each isolate at 8, 11, 14, and 18 dpi grown in three different conditions, i) 15°C, ii) 22°C, iii) 22°C in the presence of fungicide (0.05 ppm propiconazole). The same protocol described for growth assays above was also used to assess fungicide resistance. At each time point, digital images of each plate were captured and analyzed using the AIA protocol described in Lendenmann et al. (2014). We recorded measures of gray value ranging from 0 (black) to 255 (white). By using the gray value measurement from each colony on each plate and replication, we obtained the mean gray value estimate for each isolate. For a more intuitive metric of melanization, we subtracted each mean gray value from 255 to obtain a scale for melanization ranging from 0 (white) to 255 (black).

### Statistical analyses

We used model-corrected, log-transformed least-square means (LSmeans) for each host×isolate combination and for each *in planta* trait obtained by Dutta et al. (2020). The host specialization (*i.e.* the affinity of isolates for specific hosts to maximize trait performance) index for each isolate was represented by the adjusted coefficient of variation (*i.e.* logarithm of adjusted variance/mean) of LSmeans for reproduction among all 12 hosts (Döring and Reckling, 2018; Dutta et al. 2020). The LSmeans of each isolate for all the *in vitro* traits was extracted using a one-way ANOVA. To visualize trait variation among the isolates, we constructed a clustered heatmap with the R package ComplexHeatmap v.2.2.0 (Gu et al. 2016) using the *z-scores* (mean=0 and sd=1) of LSmeans for each isolate. The *z-scores* were used to reduce variability among different traits as they were expressed in different units. We excluded the morphological stress response data from the heatmap as clustering did not allow for binary data. The hierarchical clustering was performed following the agglomerative algorithm “complete” based on Euclidean distances.

We performed a genome wide association study (GWAS) for all 51 traits and 145 isolates based on 716’619 SNPs. Prior to GWAS, we conducted principal component analysis (PCA) to investigate the population structure and constructed a genetic relatedness matrix (GRM) among isolates using TASSEL v.20200220 (Bradbury et al. 2007). Both PCA and GRM were used to control the inflation of false positive SNPs due to population structure in the GWAS. The GWAS were performed by using a mixed linear model (MLM; Yu et al. 2005) including the genetic relatedness as a random factor in the R package GAPIT v.3.0 (Tang et al. 2016). PCA was excluded from the GWAS as indicated by a Bayesian information criterion (BIC) test in GAPIT for the optimal model fit. We used the stringent Bonferroni threshold (α = 0.05), false discovery rate (FDR) at 5% and 10% using the R package q-value (Storey and Tibshirani, 2003) to identify significant genome-wide SNPs.

The SNP-based heritability (*h*^*2*^_*snp*_; equivalent to narrow-sense heritability) for each trait was estimated using the GCTA tool v.1.93.0 (Yang et al. 2011). The *h*^*2*^_*snp*_ was estimated using an GREML approach including the GRM as a predictor. To investigate whether traits are under shared genetic control, we estimated genetic correlations among all traits using allelic effects across genome-wide SNPs extracted from each GWAS analysis. Phenotypic correlations among all traits were performed using the *z-scores* of the phenotypes. Both correlations were performed by using Pearson’s correlation analysis in RStudio. As the morphological stress response was recorded as a binary trait, we used point biserial correlation (Diana Kornbrot, 2014) to correlate with other quantitative phenotypes using the R package ltm v.1.1-1 (Dimitris Rizopoulos, 2006).

## Data availability

All genome sequences are available from the NCBI Sequence Read Archive (BioProject accessions PRJNA327615 and PRJNA596434).

## Author contributions

AD and DC conceived the research. AD conducted experiments, performed data analyses, and wrote the manuscript with DC. FEH and CSF provided datasets. BAM provided funding. All co-authors edited the manuscript.

## Acknowledgments

Alice Feurtey provided helpful comments on a previous version of the manuscript. This work was supported by the Swiss Federal Office for Agriculture (BLW) in the framework of the NAP-PGREL Project Nr. 627000640.

