## Supplementary Figure for "Mapping the adaptive landscape of a major agricultural pathogen reveals evolutionary constraints across heterogeneous environments"

### Supplementary information

*Dutta et al.*

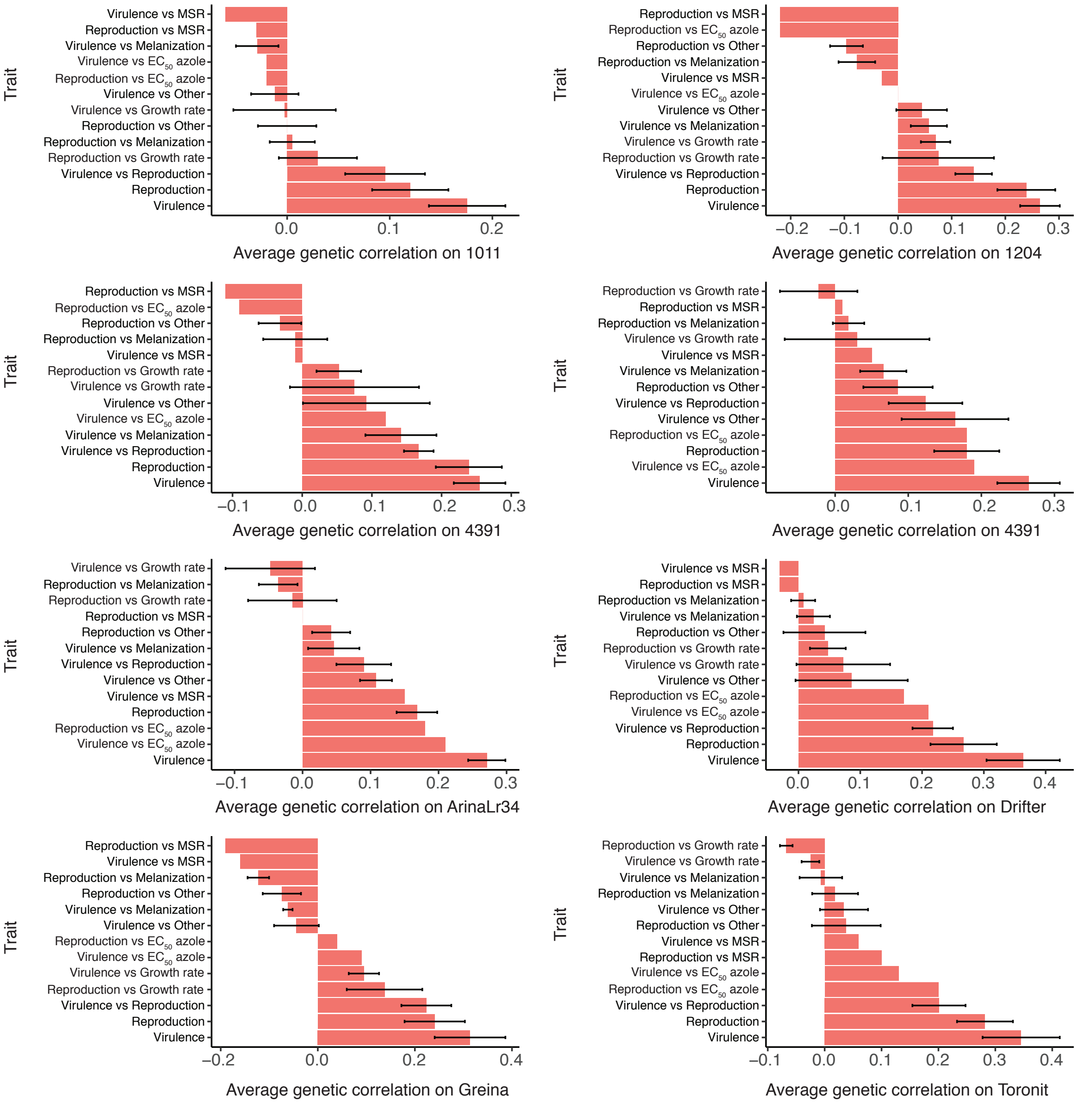


MSR= Morphological stress response (chlamydospore or hyphae formation)

**Supplementary Figure S1.** Bar plots of average genetic correlations showing the interactions of genetic control between host and non-host traits on eight hosts. Pathogen virulence (amount of necrotic lesion area) and reproduction (pycnidia density within the lesion area) were measured on 12 diverse wheat hosts. Melanization was expressed on a grayscale ranging from 0 (white) to 255 (black). Error bars indicate standard errors.

#### Supplementary Tables

(See separate Excel file)

**Supplementary Table S1.** Description of 145 *Zymoseptoria tritici* isolates with their corresponding sampling location and year used in this study.

**Supplementary Table S2.** Least-square mean values of 50 traits based on raw phenotypic data from 145 *Zymoseptoria tritici* isolates.

**Supplementary Table S3.** Summary statistics of genome-wide SNPs crossing the false discovery rate (FDR) of 10% for specific traits. SNPs are ordered according to the smallest *P-value*.

**Supplementary Table S4.** Estimates of genetic (upper diagonal) and phenotypic (lower diagonal) correlation coefficients among 50 traits in various host and non-host environments. Genetic correlations were estimated using GWAS derived allelic effects for each trait. Phenotypic correlations were estimated using standardized phenotypic values.
